# Geographic variation in the seed mycobiome of *Pseudotsuga menziesii* var. *menziesii* across the Pacific Northwest, USA

**DOI:** 10.1101/2020.11.03.367300

**Authors:** Gillian E. Bergmann, Posy E. Busby

## Abstract

Fungal symbionts occur in all plant tissues, and many aid their host plants with critical functions, including nutrient acquisition, defense against pathogens, and tolerance of abiotic stress. “Core” taxa in the plant mycobiome, defined as fungi present across individuals, populations, or time, may be particularly crucial to plant survival during the challenging seedling stage. However, studies on core seed fungi are limited to individual sampling sites, raising the question of whether core taxa exist across large geographic scales. We addressed this question using both culture-based and culture-free techniques to identify the fungi found in individual seeds collected from nine provenances across the range of Coastal Douglas-fir (*Pseudotsuga menziesii* var. *menziesii*), a foundation tree species in the Pacific Northwest and a globally important timber crop that is propagated commercially by seed. Two key findings emerged: 1) Seed mycobiome composition differed among seed provenances. 2) Despite spatial variation in the seed mycobiome, we detected four core members, none of which is a known pathogen of Douglas-fir: *Trichoderma* spp., *Hormonema macrosporum*, *Mucor plumbeus* and *Talaromyces rugulosus*. Our results support the concept of a core seed microbiome, yet additional work is needed to determine the functional consequences of core taxa for seedling germination, growth, survival and competition.

## INTRODUCTION

Fungal symbionts occur commonly in plants and impact essential plant functions such as growth and defense (Rodriguez et al. 2009). However, because the plant mycobiome (i.e. fungi in the microbiome) varies at different spatial scales (organs, individuals, sites), it is crucial to evaluate why these communities vary and whether this variation contributes to variation in plant phenotypes. As part of analyzing community variation across space, identifying fungi that are consistently present across individuals and populations, aka “core” species (Shade and Handelsman 2012), can help researchers and farmers alike focus on stable taxa that are most likely to impact host phenotypes (Busby et al. 2017).

Studies in both humans and plants alike identify core taxa as those that are present across a certain threshold of all samples analyzed (usually 80-90%, e.g. Turnbaugh et al. 2009, Huse et al. 2012, Eyre et al. 2019, Chen et al. 2020). However, this definition doesn’t account for the relative abundances of core taxa or their contribution to community similarity across samples. Alternatively, core taxa have also been identified by their relative abundances across samples (e.g. Edwards et al. 2015, Timm et al. 2018), which is often biased towards dominant taxa (Shade and Handelsman 2012). To combat these limitations, Shade and Stopnisek (2019) proposed using abundance-occupancy distributions to identify a core, an approach that ranks taxa based on their presence and abundance across groups of samples and identifies core taxa on their contribution to community composition. However, few studies have used this approach to identify core taxa (e.g. Grady et al. 2019, Stopnisek and Shade 2019), and the core mycobiome has yet to be defined for most plant tissues.

Identifying core taxa in the seed mycobiome is of particular interest since some seed fungi are vertically transmitted (Barret et al. 2015), potentially impacting plant phenotypes over multiple generations. Additionally, as the first source of inoculum in a plant’s life cycle, seed fungi may also influence seedling mycobiome assembly via priority effects (Fukami 2015), and contribute significantly to seedling survival (Nelson 2018). Due to small seed sizes, chemical defenses (Stamp 2003), and exclusionary interactions between taxa (Raghavendra et al. 2013), fungal communities inside seeds are expected to have limited membership. In such depauperate communities, the presence and function of core taxa is likely to be especially important. However, the seed mycobiome often receives less attention in comparison to foliage, roots, stems. Much of the research that has been done on the seed mycobiome has focused on seed-borne pathogens, and only a few studies have identified core fungi in crop seeds collected from single sampling sites (e.g. Rezki et al. 2018, Eyre et al. 2019, Chen et al. 2020). As such, we have limited knowledge on the overall composition and core taxa of seed mycobiomes for most plants across large spatial scales.

In this study, we used both culture-based and culture-free molecular methods to survey the mycobiome of individual Coastal Douglas-fir (*P. menziesii* var. *menziesii*) seeds in the Pacific Northwest, USA. In both culture-based and culture-free approaches we surface sterilized seeds prior to isolating microbes from the seeds, thus our study characterizes seeds present *in* rather than *on* seeds. We used our survey data to evaluate how seed mycobiome composition varies across individual seeds and seed provenances (regions of origin). We used the abundance-occupancy distribution approach to examine if the *P. menziesii* var. *menziesii* seed mycobiome had consistent, or “core,” taxa across seed provenances. We then used our core mycobiome data to evaluate how core composition varies across seed provenances. Finally, we opportunistically sampled the mycobiome of a small number of seeds from New Zealand-grown Douglas-fir for qualitative comparisons to the US samples.

## MATERIALS AND METHODS

### Study system

We conducted our seed mycobiome survey in Coastal Douglas-fir (*Pseudotsuga menziesii* var. *menziesii*), a foundational member of coniferous forests in western North America (Uchytil 1991) and a major timber crop around the world (Eilmann et al. 2013, Watts et al. 2015). While a number of studies have examined seed fungi in this species, they rely on morphological identification (Salisbury 1955, Bloomberg 1966), focus on seeds from nurseries (Bloomberg 1966, Morgenstern et al. 2014), or focus on the presence of pathogens like *Fusarium* (James et al. 1989, Hoefnagels and Linderman 1999) and *Rhabdocline* (Morgenstern et al. 2014). Thus, the complete seed mycobiome composition of *P. menziesii* var. *menziesii* across the species’ ranges is unknown.

### Seed collection and preparation

To assess the association between seed mycobiome composition and seed provenance, we sampled seeds across the tree species’ native range. Seeds were collected from eight provenances in the western United States by Silvaseed Company Foresters (Washington, USA). The provenances represent seed sources in wild forests spread across a latitudinal gradient from Northern Washington to Central California (TABLE 1). Fresh cones were harvested from the ground in wild forests within a five to ten-mile radius at each zone, within elevations of 0 to 1500 feet (Gerdes, personal communication). Cones were checked for recent pollination prior to harvest, and seeds were prepared for storage through drying, removal from cones, dewinging and removal of debris (Gerdes, personal communication). Processed seeds were placed in cold storage and kept there for several months to several years before being provided for this study (Gerdes, personal communication).

**TABLE 1.**
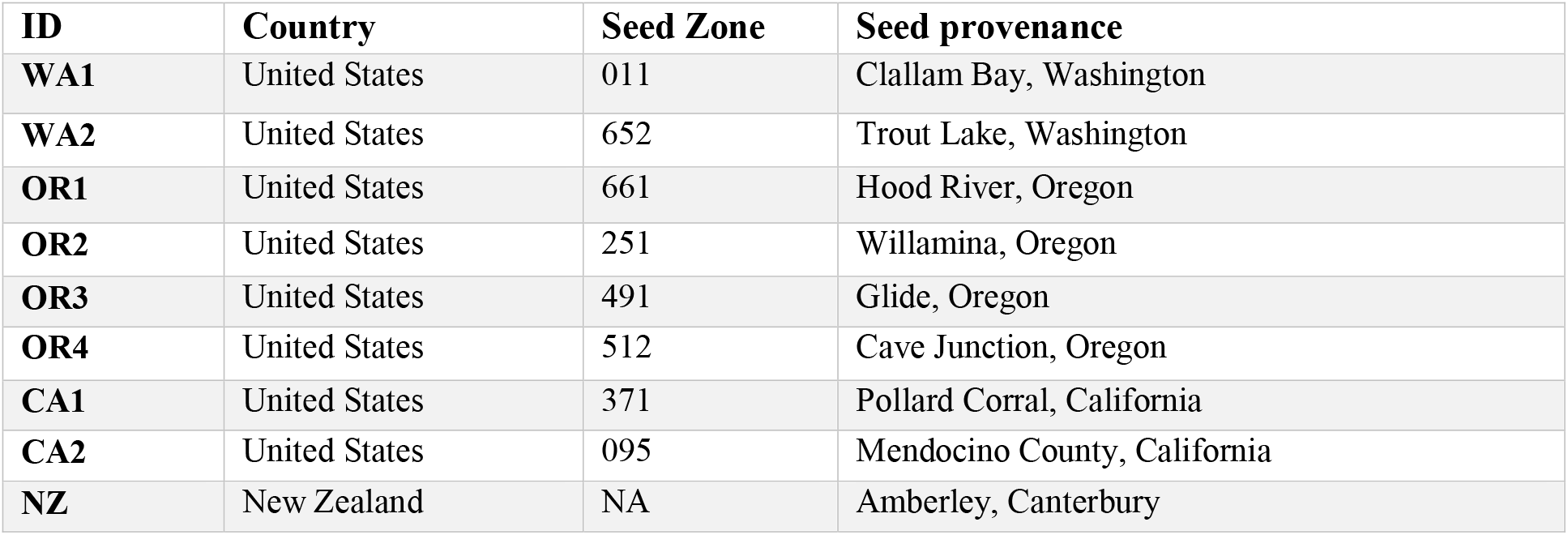
Summary of seed provenances collected for characterization. Seed zones reported by Silvaseed Company Foresters (Washington, USA).

Another provenance was in New Zealand (TABLE 1), outside of the tree’s native range, where seeds were collected opportunistically to make qualitative comparisons against the United States provenances. For sampling the introduced provenance of *P. menziesii* var. *menziesii* in New Zealand, we used seeds from three lines grown in an orchard in Amberley, Canterbury collected by Proseed New Zealand (Canterbury, NZ; TABLE 1). The seed lines sampled represented populations that were introduced to New Zealand in 1959 from 44 provenances across Washington, Oregon, and California, and were grown in the orchard after introduction (<u>Proseed NZ 2019; van Ballekom, personal communication</u>). Seeds were harvested in 2011 from cones located on branches 2.5-3 meters above the ground (van Ballekom, personal communication). Following harvest, cones were air and kiln-dried, and seeds were then extracted and stored at 4((van Ballekom, personal communication). Seeds were assessed for viability every two years following harvest (van Ballekom, personal communication).

### Culture-based characterization

We first used culture isolations and Sanger sequencing to identify the culturable mycobiome of *P. menziesii* var. m*enziesii* seeds. First, we stratified all seeds at 4°C for four weeks following the protocol developed by Mujic (2015) to promote endophyte emergence. We sterilized 100 individual seeds from each United States provenance using ethanol (95%) with TWEEN 80 (0.01%) for 10 seconds, hydrogen peroxide (30%) for 2 minutes, ethanol (70%) for 2 minutes and sterilized deionized water for 1 minute. Seeds were gently agitated in each solution and allowed to dry for ten minutes following the soak in sterilized distilled water. All seeds were plated onto full strength potato dextrose agar (PDA) and incubated at room temperature in ambient light for up to 3 weeks, and emerging endophytes were sub-cultured onto fresh media. Sterilization of seed samples was confirmed by rolling seeds on the media used for endophyte isolation, removing seeds from the media, then monitoring for microbial growth over a 10-day period. A total of 100 seeds from each New Zealand line were prepared in a similar manner with the following differences: seeds were sterilized in sodium hypochlorite (0.5%) instead of hydrogen peroxide, they were cut in half before plating on malt yeast extract agar (MYE) agar amended with the antibiotics chlortetracyline hydrochloride (50mg/L) and streptomycin sulphate (250 mg/L), and sterilized distilled water was also plated to check for contamination.

We extracted genomic DNA from all United States isolates using the Sigma Extract-n-Amp kit (Sigma-Aldrich, Missouri, USA). ITS1-F (Gardes and Bruns 1993) and LR3 (Vilgalys 2018) primers were used to amplify the Internal Transcribed Spacer (ITS) region via PCR (Schoch et al. 2012). PCR reactions had a total volume of 25uL and included 2uL of genomic DNA. A Chelex 100 buffer (5%, Bio-Rad Laboratories) method was used to extract DNA from New Zealand isolates (Hill et al., 2017), and ITS4 and ITS5 (White et al. 1990) primers were used to amplify the ITS2 region in 50uL PCR reactions with 2uL of genomic DNA. All PCR amplicons were visualized using gel electrophoresis. United States amplicons with a visible band were sent to MCLAB (San Francisco, CA) for Sanger sequencing, and New Zealand amplicons with a visible band were Sanger sequenced at the Bio-Protection Research Centre. All raw sequence reads were cleaned with SeqTrace (Stucky 2012), and the final reads were used to estimate taxonomic identity with query results on the UNITE (Kõljalg et al. 2005; http://unite.ut.ee/index.php) database. Identified sequences were submitted to GenBank (accessions MT663418-MT663509).

### Analysis of culture-based data

For the United States samples, we conducted chi-square tests to evaluate association between seed provenance and cultured fungal frequency, for both total endophyte frequency, and frequency of any common taxa. Frequency was defined as the number of times an endophyte was isolated out of the total number of seeds cultured per provenance. Chi-squared tests were performed in R 3.6.3 (R Core Team 2020) using the MOSAIC (Pruim et al. 2019) package. We qualitatively compared the fungal communities between the United States and New Zealand for similarities and differences in the taxa identified, with isolates from the New Zealand lines combined to represent one provenance.

### Preparation of the Illumina MiSeq library

In addition to the culture-based data, we generated culture-independent data for the same batch of seeds to identify culturable and unculturable members of the seed mycobiome. A total of 180 stratified seeds (20 per provenance) from the same seed lots were surface sterilized in sodium hypochlorite (.5%) with TWEEN (0.01%) for 1 minute, followed by three washes in sterile water for 1 minute each. Seeds from the New Zealand lines were combined as one provenance for sampling. After drying for ten minutes, seeds were cut in half with a sterile scalpel and tissue from each individual was homogenized in a Geno/Grinder 2010 tissue homogenizer (Spex SamplePrep, New Jersey, USA). Total genomic DNA was extracted from the samples using a 96 Well Synergy Plant DNA Extraction Kit (OPS Diagnostics LLC, New Jersey, USA) according to the manufacturer’s protocol. The four corners of each extraction plate were processed without seeds to serve as extraction blanks in the library preparation process. Following extraction, samples were purified with Agencourt AMPure XP beads (beads to DNA, 1 to 1 ratio; Beckman Coulter Inc., Pasadena, CA, USA).

The amplicon library was constructed following two-stage PCR amplification. Stage one PCR was performed using ITS1-F and ITS2 primers (Toju et al. 2012) to target the ITS1 region. Forward and reverse primers were modified with adaptors and 3-6bp long heterogeneous spacers. PCR reactions were 25uL in volume, and consisted of 12.5uL MyFi mix polymerase (Bioline, Tennessee, USA), 1uL of each primer and 5 uL of genomic DNA. Each plate had three extraction blanks and a PCR negative control. Stage one PCR conditions were as follows: 95(for 3 minutes, 34 cycles of 95(30s), 50(30s) and 72(15s), with a final extension step for 5 minutes at 72.(Stage two PCR reactions were 20uL in volume, with 2 uL of amplicon and primers containing Illumina adaptors and indexes (Hamady et al. 2008). Extraction blanks and PCR controls were carried over from the first stage. Stage two PCR conditions were: 95(for 3 minutes, 8 cycles of 95(30s), 55(30s) and 72(30s), with a final extension step of 72(for 2 minutes. All amplicons were purified and normalized using Just-a-Plate purification and normalization plates (Charm Biotech, Missouri, USA), and quantified with Qubit (ThermoFisher Scientific, Massachusetts, USA). Equimolar concentrations of each sample were pooled and sent to the Oregon State University Center for Genome Research and Biocomputing for 300-bp, paired-end sequencing on the Illumina MiSeq platform. The raw sequences were submitted to the NCBI Small Read Archive (accession PRJNA649965).

### Processing of Illumina MiSeq library

We used the following method to prepare our MiSeq library for analysis. Sequences were demultiplexed using Pheniqs (Galanti et al. 2017) with a maximum likelihood confidence threshold of 0.995. Adaptors and the 5’ gene primer were trimmed using Cutadapt 2.10 (Martin 2011). Adaptors and the 3’ gene primer were trimmed using SeqPurge (Sturm et al. 2016). Sequences were truncated using Cutadapt to a final length of 230bp for forward and reverse reads based on sequence quality reports. After trimming, host sequences were removed with VSEARCH (Rognes et al. 2016) against the ITS region of *P. menziesii* (Gernandt and Liston 1999). Sequences were filtered at a maximum expected error rate of 2bp, and DADA2 (Callahan et al. 2016) was used to denoise and remove chimeras from the filtered reads. Taxonomy was assigned using the UNITE eukaryote database (Kõljalg et al. 2005; https://doi.org/10.15156/BIO/786371). The denoised data was combined into a single phyloseq object (McMurdie and Holmes 2013) that was used in downstream processing and analysis. Taxa not assigned to the kingdom Fungi were removed, and remaining taxa were re-assigned with the UNITE fungal database (Kõljalg et al. 2005; https://doi.org/10.15156/BIO/786369). Controls and putative contaminants (taxa with more than 2% of reads in the controls) were removed, as well as samples with a sequencing depth below 2000 reads. Filtered reads were clustered into operational taxonomic units (OTUs) with DECIPHER (Wright 2016) at a similarity threshold of 97%. Separate phyloseq objects were created with all OTUs included (full OTU), singletons removed and taxa present in less than 2.5% of samples removed (2.5% cutoff). OTU reads were rarefied to proportional abundance for all three phyloseq objects.

### Analysis of fungal community composition using culture-free data

We used multiple analyses to compare, contrast and visualize the composition of the sequence-based mycobiome and test its association with seed provenance. We visualized the community composition of the mycobiome for individual seeds in each provenance using a non-metric multidimensional scaling (NMDS) ordination plot (Bray-Curtis distances for all samples) of the 2.5% cutoff phyloseq object. We performed permutational multivariate analysis of variance (PerMANOVA) to test the association between seed provenance and community composition in the United States. OTU, spatial (latitude) and environmental (Mean Annual Precipitation) vectors were calculated to identify potential drivers of mycobiome composition across seed provenances. Shannon diversity was calculated for all samples using the full OTU phyloseq object, and a one-way analysis of variance (ANOVA) test was conducted on Shannon diversity across seed provenances. All analyses were performed in R 3.6.3 (R Core Team 2020) using the PHYLOSEQ (McMurdie and Holmes 2013), VEGAN (Okansen et al. 2019), and GGPLOT2 (Wickham 2016) packages.

### Identification and analysis of the core mycobiome using culture-independent data

We used an abundance-occupancy distribution approach to identify core taxa in the United States seeds. This identification process was performed in R 3.6.3 (R Core Team 2020) using code modified from Shade and Stopnisek (2019). Briefly, we calculated the occupancy (percent detection in each provenance) and relative abundance (percent of total reads in each sample) of all OTUs in the Illumina MiSeq library. We then used these occupancy and abundance values to calculate an index for each OTU. We ranked OTUs by their abundance-occupancy indices, and calculated the contributions of each OTU to the Bray-Curtis (BC) similarity of the total mycobiome across samples. We plotted the percent BC similarity as a function of the contributions of these ranked OTUs, and identified core taxa using the elbow method, a conservative approach based on 1^st^ order differences in the slope of the BC similarity curve between OTUs. We created a separate phyloseq object of these core taxa for downstream analyses, and checked if these taxa included known pathogens of *P. menziesii* var. *menziesii* in the U.S. National Fungus Collections Fungus-Host Database (https://nt.ars-grin.gov/fungaldatabases/fungushost/fungushost.cfm). We identified the percentage of samples where core taxa were members, and visualized the relative abundance of these taxa across seeds. We then performed a PerMANOVA test to examine the association between seed provenance and core composition.

## RESULTS

### Taxonomic composition of the culture-based mycobiome

We used both culture-based and culture-independent approaches to survey the mycobiome of individual *P. menziesii* var. *menziesii* seeds across nine provenances in the United States and New Zealand. In the culture-based effort, we isolated a total of 93 fungal strains from 1100 seeds across all provenances (8.9% isolation frequency). All seeds had either zero or one fungus, and fungal isolation frequency per provenance ranged from 0 to 28%. We identified all isolates to the phylum *Ascomycota* representing 4 classes, 6 families, 7 genera and 13 species (TABLE 2). The most common taxa in the United States were *Trichoderma* and *Hormonema macrosporum*, representing 35.56% and 26.67% of the isolates respectively (TABLE 2). The most common taxon in New Zealand was *Hormonema dematioides*, representing 93.75% of the isolates (TABLE 2).

**TABLE 2.** Fungal taxa isolated and identified using Sanger sequencing in order of relative abundance, including information on seed provenance, relative abundance by country, and accession number.

| Species | Source provenance | US rel. abund. (%) | NZ rel. abund. (%) | Representative Accession |
| --- | --- | --- | --- | --- |
| *Hormonema dematioides* | NZ | 0 | 93.75 | MT663448 |
| *Hormonema macrosporum* | CA1 | 35.56 | 0 | MT663419 |
| *Trichoderma koningii* | WA1, OR1, CA1 | 26.67 | 0 | MT663431 |
| *Trichoderma citrinoviride* | CA1 | 8.89 | 0 | MT663418 |
| *Trichoderma spp.* | WA1 | 6.67 | 0 | MT663505 |
| *Unknown Tympanidaceae spp.* | CA1 | 6.67 | 0 | MT663433 |
| *Mycopappus spp.* | OR1 | 4.44 | 0 | MT663497 |
| *Aspergillus pseudoglaucus* | NZ | 0 | 4.17 | MT663450 |
| *Trichoderma ovalisporum* | WA1 | 2.22 | 0 | MT663509 |
| *Trichoderma stellatum* | WA1 | 2.22 | 0 | MT663508 |
| *Trichothecium roseum* | CA2 | 2.22 | 0 | MT663447 |
| *Trichoderma harzianum* | NZ | 0 | 2.08 | MT663452 |
| **Total** |  | 100 | 100 |  |

### Taxonomic composition of the culture-free mycobiome

We obtained a total of 3,453,150 sequences from 170 individual seeds across nine provenances in our MiSeq library, with between 2,024 and 42,794 sequences per sample. Our final dataset had 294 Operational Taxonomic Units (OTUs) representing 3 phyla (*Ascomycota, Basidiomycota, Mucoromycota*), 14 classes, 32 orders, 51 families and 63 genera (FIG. 1). Individual seeds had between 1 and 19 OTUs, and some seeds were dominated by a single taxon while others included several (FIG. 1). The five most common species across all provenances were *Trichoderma spp.* (20.7% of reads)*, Hormonema macrosporum* (9.59%)*, Mucor plumbeus* (7.09%)*, Talaromyces rugulosus* (6.52%) and *Uncobasidium spp.* (5.18%).

**Figure 1.**
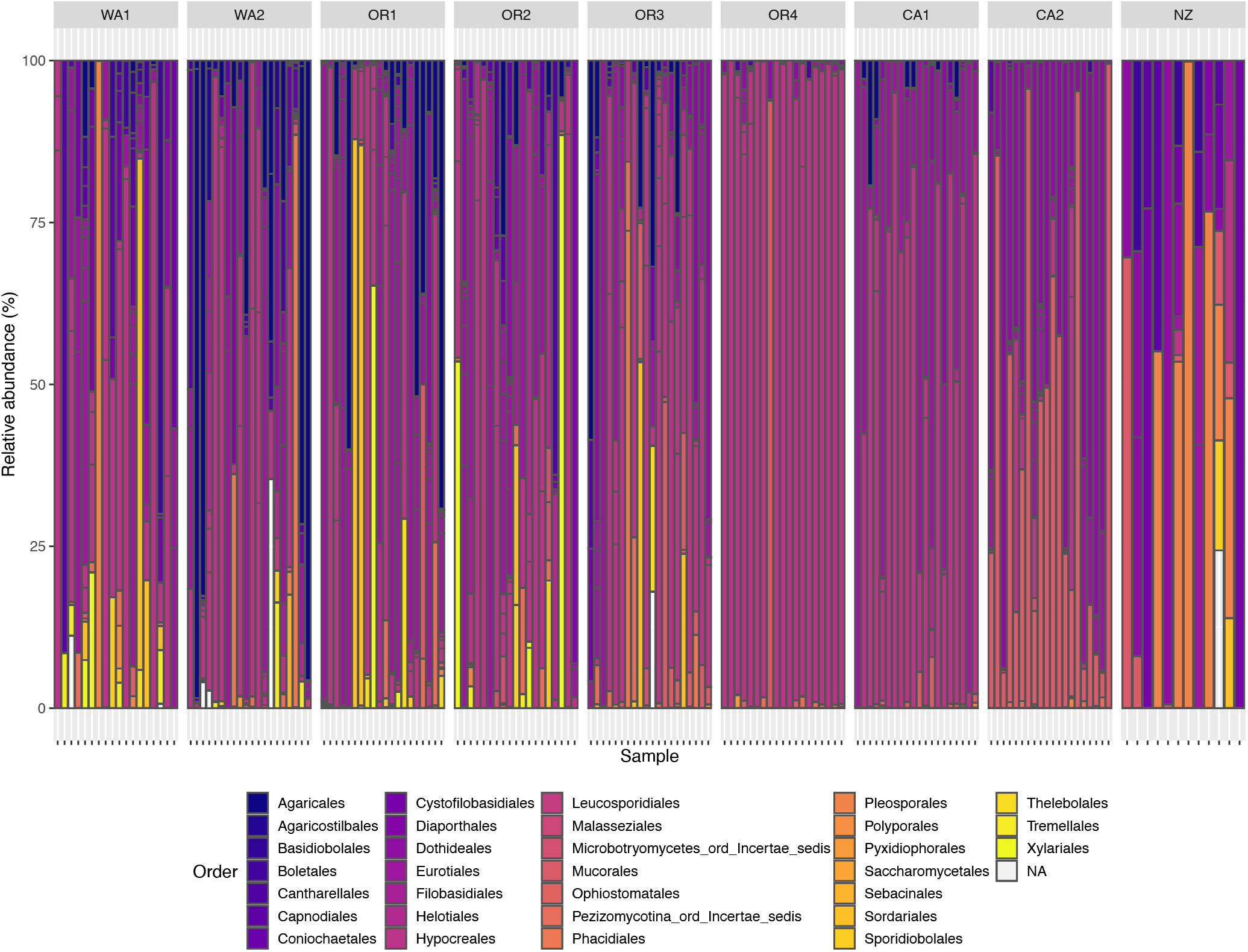
Taxonomic distribution of the culture-free seed mycobiome across all samples (along horizontal axis). Samples are organized into seed provenances by latitude. Each color in sample bars represents a fungal order, and lines along the bars separate individual OTUs.

### Provenance-associated variation in mycobiome frequency and composition

In both our culture-based and sequence-based approaches, seed mycobiome composition differed significantly among United States provenances. Culturable fungal frequency (X^2^ = 108.46, p <2.2×10^−16^), culturable *Trichoderma* frequency (X^2^ = 44.801, p = 1.495×10^−7^) and culturable *Hormonema macrosporum* frequency (X^2^ = 128.9, p <2.2×10^−16^) differed among United States provenances. In the sequence-based approach, seed provenance explained 30.9% of the variation in mycobiome composition (PerMANOVA; df =7, F=9.4575, p=0.001; FIG. 2A), however, beta-dispersion differed among provenances. Shannon diversity also varied significantly between seed provenances (ANOVA; df = 7, F=12.3, p=2.3×10^−12^).Variation in culture-free mycobiome composition among United States provenances was correlated with latitude (vector analysis; r=0.546, p=0.001), and the abundance of *Trichoderma spp.* (vector analysis; r = 0.412, p = 0.001), *Hormonema macrosporum* (vector analysis; r = 0.239, p =0.001) and *Mucor plumbeus* (vector analysis; r = 0.235, p = 0.001; FIG. 2B).

**Figure 2.**
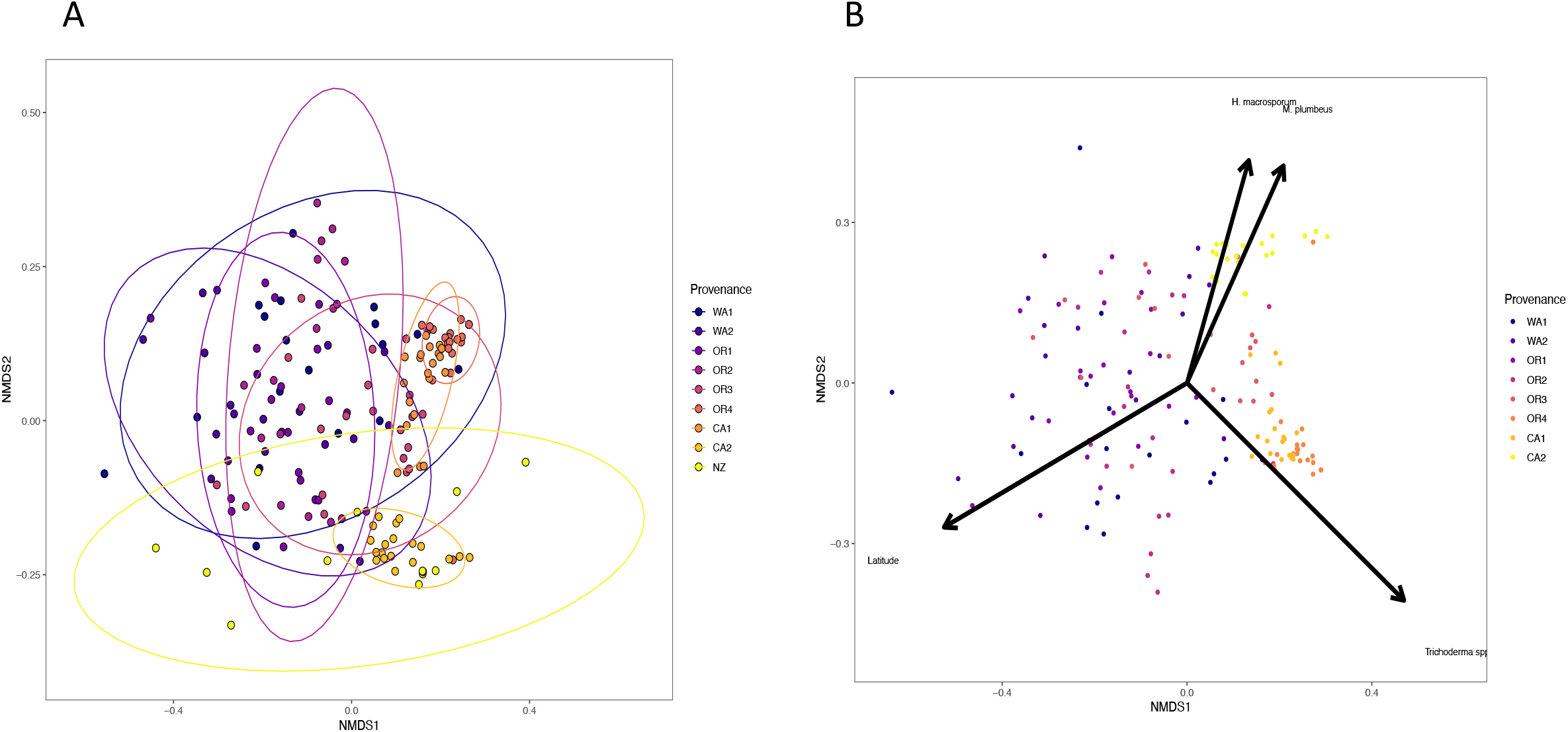
(A) Non-metric dimensional scaling (NMDS) plot of culture-free seed mycobiome composition using a Bray-Curtis dissimilarity matrix. Each point represents a single seed sample (n=170), and each color represents a seed provenance. (B) Non-metric dimensional scaling (NMDS) plot of the culture-free seed mycobiome from the United States provenances. Each point represents a single seed (n=156), and each color represents a seed provenance. Environmental and OTU vectors with significant association to community composition are shown (latitude: r=0.546, p=0.001; *Trichoderma spp*.: r = 0.412, p = 0.001; *Hormonema macrosporum*: r = 0.239, p =0.001; *Mucor plumbeus*: r = 0.235, p = 0.001).

We also observed differences in mycobiome composition between the United States and New Zealand. *Trichoderma* was the only cultured fungal genus found in both the US and New Zealand, and species richness was lower in New Zealand than in the United States for cultured fungi. Culture-free community composition differed significantly between New Zealand seeds and seeds from all United States provenances (FIG. 2A).

### *Core mycobiome of P. menziesii var. menziesii* seeds

We identified core taxa of United States seeds from the culture-independent data using the elbow method on an occupancy-abundance distribution (Shade and Stopnisek 2019). Four taxa were identified as core with this method: *Trichoderma spp.* (OTU.1)*, Hormonema macrosporum* (OTU.2)*, Mucor plumbeus* (OTU.4) and *Talaromyces rugulosus* (OTU.3). These core taxa represented 43.9% of the total fungal reads, and were present in 38.7%, 50%, 48.2% and 44.6% of seeds respectively. Relative abundance of these taxa was variable across seeds where they were present, ranging from less than 1% to 100% of reads per individual (FIG. 3). Seed provenance explained 33.9% of the variation in core mycobiome composition (PerMANOVA; df =7, F=10.236, p=0.001).

**Figure 3.**
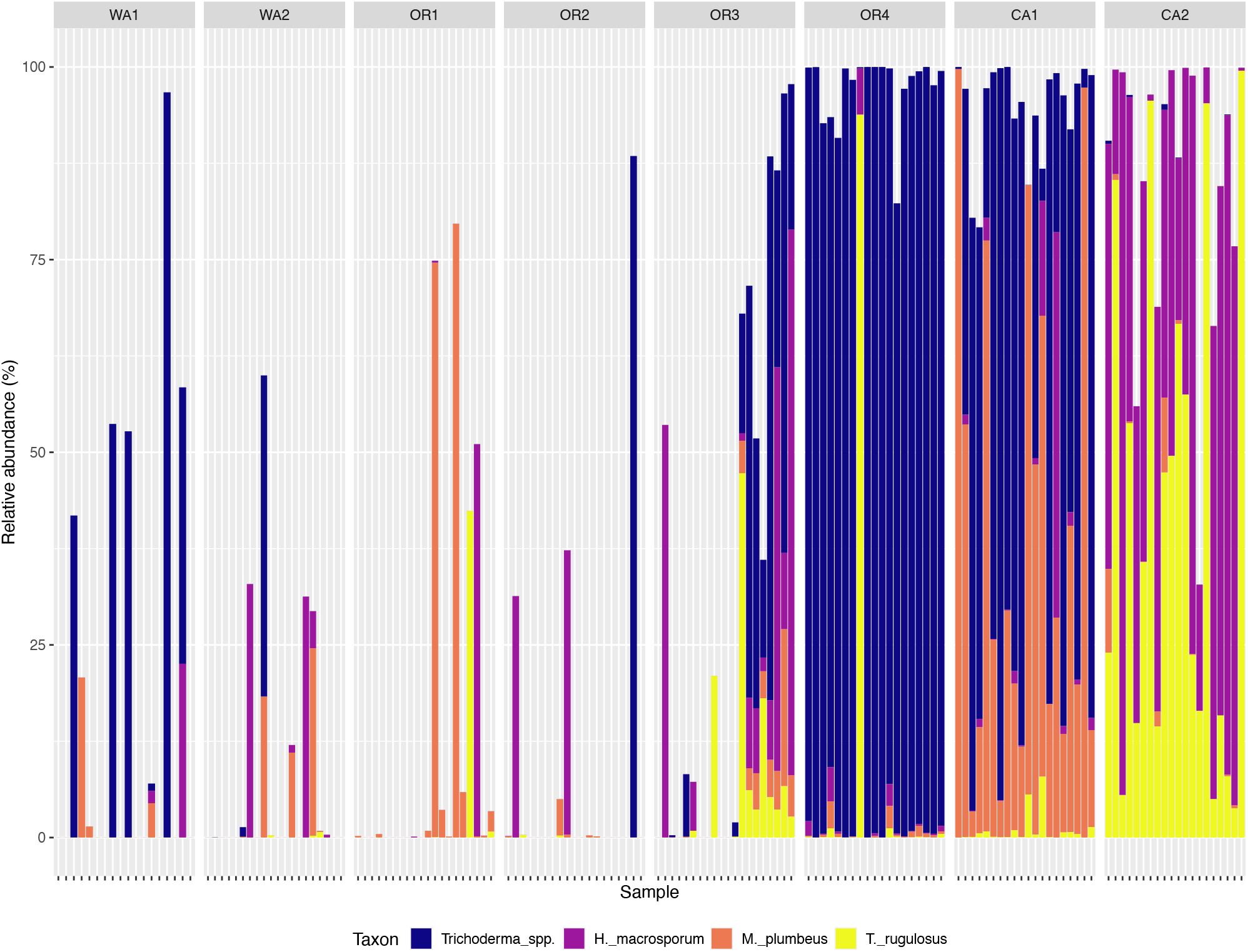
Core mycobiome composition of United States seeds defined by the elbow method on an abundance-occupancy distribution. Each bar represents an individual seed sample, and samples are grouped in provenances by latitude. Bar colors represent each core taxon, and bar heights represent the total relative abundance of the core taxa.

## DISCUSSION

We found that individual seeds of *P. menziesii* var. *menziesii* from the United States and New Zealand contain predominantly Ascomycetes in the classes Sordariomycetes (29.5% of taxa) and Eurotiomycetes (20.5% of taxa). Five genera were present in both the culture-based and culture-independent approaches: *Aspergillus, Hormonema, Mycopappus, Trichoderma* and *Trichothecium*. The majority of identified taxa are not known pathogens of *P. menziesii var. menziesii*, which include *Rhabdocline* and *Nothophaeocryptopus gaeumanii* (Sherwood-Pike et al. 1986, Morgenstern et al. 2014, Gómez-Gallego et al. 2019).

The communities of individual seeds contained few fungal OTUs, and were often dominated by a single OTU. Additionally, seeds either had zero or one culturable fungus. This pattern of cultured fungal frequency supports the Primary Symbiont Hypothesis, which states that seeds have a dominant microbe that is functionally important (Newcombe et al. 2018). The culture-independent data, however, only provides limited support for this hypothesis, since seeds from some provenances were dominated by a single OTU while others were not. This suggests that there are unculturable and/or inviable fungi present in seeds that are not accounted for in culture-based surveys. Our surface sterilization protocol may have affected the viability of some seed fungi. Future studies pairing culture-based and culture-free mycobiome identification across a broader range of plants will be better poised to validate or refute whether a primary symbiont exists, and what factors contribute to its presence.

We found that there was a strong association between seed provenance and fungal community composition in both the culture-based and culture-independent data. Across the United States provenances, community dissimilarity was correlated with latitude, but not mean annual precipitation. Distance and other abiotic environmental factors may thus be influential for fungal community composition in *P. menziesii* var. *menziesii* seeds. Environmental and spatial distance are both important factors in mycobiome variation (Barge et al. 2019), and local site conditions have previously been associated with seed mycobiome composition (Klaedtke et al. 2016, Rezki et al. 2018). The relative influence of environmental heterogeneity, dispersal and local processes on community composition is the basis of metacommunity theory (Leibold and Chase 2018), which could be a useful framework for studying mycobiomes. Because we have limited metadata on our seed provenances, however, we cannot say which spatial or environmental factors are influencing the variation we observed. Additionally, the variation could be confounded by differences in storage conditions, and how the seeds were sampled (ground vs. branch, wild forest vs. plantation). Future work should target the seed mycobiome along specific environmental and spatial gradients to determine how environmental filtering and dispersal impact community composition across such large spatial scales.

We observed differences in the fungal communities between the United States and New Zealand. Cultured species richness and uncultured Shannon diversity were lower in the New Zealand seeds than in the United States, and seed communities in New Zealand were distinct from those in the United States (TABLE 2, FIG. 2A). These differences could be due to distinct species pools in each country and limited migration of seed fungi when *P. menziesii* var. *menziesii* was introduced to New Zealand. However, differences in the sampling and culture-based identification methods confound the variation we observed between countries, so we cannot say what is driving the variation. Still, the New Zealand seeds represent an interesting outgroup for comparison against seeds within *P. menziesii* var. *menziesii’s* native range.

In addition to identifying an association with provenance, we found that United States seeds have a core mycobiome of four common taxa across provenances. These core taxa were identified based on their presence across multiple seeds and seed provenances and their contribution to the similarity across all communities sampled. This approach allowed us to identify core taxa that may be ecologically important in terms of their contribution to mycobiome composition and function. However, these taxa would not be identified as core with the membership or abundance methods, since they were not present in more than 50% of seeds and their relative abundance was highly variable between provenances. This highlights the importance of explicitly defining what is meant by “core” mycobiome. Explicitly stating the scale of inference is also important when identifying a core mycobiome, as the presence and size of a core changes with spatial scale (Vandenkoornhuyse et al. 2015). The variability we observed in core composition was also associated with seed provenance, suggesting that environmental filters and dispersal between provenances may also determine if a core establishes. Since we isolated several of these core taxa in culture, future studies in this system can test the impacts of different processes on core assembly, and determine if these taxa are functionally important.

There is little overlap between the seed mycobiome identified in this study and the taxa found in Douglas-fir needles and roots in previous studies. We did not find the four major foliar pathogens (*Rhabdocline* and *Nothophaeocryptopus gaeumannii;* Sherwood-Pike et al. 1986, Morgenstern et al. 2014, Gómez-Gallego et al. 2019) of *P. menziesii* var. *menziesii* in seed, and 3.1% and 3.7% of the culture-free OTUs have been observed in needles (Carroll and Carroll 1978, Daniels et al. 2018) and roots (Hoff et al. 2004) respectively. Plant compartment is a crucial factor in structuring the mycobiome (Coleman-Derr et al. 2016, Cregger et al. 2018), and the tissue specificity of *P. menziesii* var. *menziesii* seeds could be due to their unique structure and chemical defenses. Alternatively, different dispersal rates of fungi between seeds, other tissues and the ambient environment could contribute to the observed composition differences. Since we surveyed the mycobiome composition of seeds from an indeterminate number of trees, we cannot discern the relative effects of these metacommunity processes on community composition. Future work treating the full mycobiome of *P. menziesii* var. *menziesii* as a metacommunity could identify the contributions of these environmental filters and dispersal rates across plant tissues to community assembly and composition.

## CONCLUSIONS

We found that the seed mycobiome of *P. menziesii* var. *menziesii* for the United States and New Zealand varies across geographic scales and contains a putative core across provenances. The variation we observed across seeds, tissues and provenances emphasizes the need to study mycobiomes as metacommunities across multiple scales. While we found that among-provenance heterogeneity in the seed mycobiome was tied to the latitudinal gradient of provenances studied, the specific environmental and spatial factors structuring the mycobiome are still unknown. Future work using the cultured mycobiome isolated here could test how metacommunity processes affect total and core mycobiome assembly, and if the mycobiome influences seedling health.

## ACKNOWLEDGEMENTS

We thank Edward Barge, Devin Leopold, Javier Tabima and Shikhar Hatwal for their assistance with laboratory work and bioinformatics. We thank Helen Whelan, John Hampton and Travis Glare at the Bio-Protection Research Centre for their mentorship during the culture-based work in New Zealand. Special thanks go to Kyle Gervers for his assistance on data analysis and manuscript revisions. We thank Rachel Vannette, Marshall McMunn, Johan Leveau and Tyler Laird for their feedback on the manuscript.

## Footnotes

1 GE Bergmann is now affiliated with the Graduate Group in Ecology at the University of California-Davis.

2 All data and scripts are available at Dryad DOI: https://doi.org/10.25338/B8HS6B.

## REFERENCES

Barge EG, Leopold DR, Peay KG, Newcombe G, Busby PE. 2019. Differentiating spatial from environmental effects on foliar fungal communities of Populus trichocarpa. J Biogeogr. 0, doi:10.1111/jbi.13641.

Barret M, Briand M, Bonneau S, Préveaux A, Valière S, Bouchez O, Hunault G, Simoneau P, Jacques M-A. 2015. Emergence Shapes the Structure of the Seed Microbiota. Appl Environ Microbiol. 81:1257–1266, doi:10.1128/AEM.03722-14.

Bloomberg WJ. 1966. THE OCCURRENCE OF ENDOPHYTIC FUNGI IN DOUGLAS-FIR SEEDLINGS AND SEED. Can J Bot. 44:413–420, doi:10.1139/b66-050.

Busby PE, Soman C, Wagner MR, Friesen ML, Kremer J, Bennett A, Morsy M, Eisen JA, Leach JE, Dangl JL. 2017. Research priorities for harnessing plant microbiomes in sustainable agriculture. PLOS Biol. 15:e2001793, doi:10.1371/journal.pbio.2001793.

Callahan BJ, McMurdie PJ, Rosen MJ, Han AW, Johnson AJA, Holmes SP. 2016. DADA2: High-resolution sample inference from Illumina amplicon data. Nat Methods. 13:581–583, doi:10.1038/nmeth.3869.

Carroll GC, Carroll FE. 1978. Studies on the incidence of coniferous needle endophytes in the Pacific Northwest. Can J Bot. 56:3034–3043, doi:10.1139/b78-367.

Chen X, Krug L, Yang H, Li H, Yang M, Berg G, Cernava T. 2020. Nicotiana tabacum seed endophytic communities share a common core structure and genotype-specific signatures in diverging cultivars. Comput Struct Biotechnol J. 18:287–295, doi:10.1016/j.csbj.2020.01.004.

Coleman-Derr D, Desgarennes D, Fonseca-Garcia C, Gross S, Clingenpeel S, Woyke T, North G, Visel A, Partida-Martinez LP, Tringe SG. 2016. Plant compartment and biogeography affect microbiome composition in cultivated and native Agave species. New Phytol. 209:798–811, doi:10.1111/nph.13697.

Cregger MA, Veach AM, Yang ZK, Crouch MJ, Vilgalys R, Tuskan GA, Schadt CW. 2018. The Populus holobiont: dissecting the effects of plant niches and genotype on the microbiome. Microbiome. 6:31, doi:10.1186/s40168-018-0413-8.

Daniels HA, Cappellazzi J, Kiser JD. 2018. Microbiome Diversity of Endophytic Fungi across Latitudinal Gradients in West Coast Douglas-Fir (Pseudotsuga menziesii) Foliage. J Biodivers Manage For. 7, doi:10.4172/2327-4417.1000203.

Edwards J, Johnson C, Santos-Medellín C, Lurie E, Podishetty NK, Bhatnagar S, Eisen JA, Sundaresan V. 2015. Structure, variation, and assembly of the root-associated microbiomes of rice. Proc Natl Acad Sci. 112:E911–E920, doi:10.1073/pnas.1414592112.

Eilmann B, Vries SMG de, Ouden J den, Mohren GMJ, Sauren P, Sass-Klaassen U. 2013. Origin matters! Difference in drought tolerance and productivity of coastal Douglas-fir (Pseudotsuga menziesii (Mirb.)) provenances. For Ecol Manag. 302:133–143, doi:10.1016/j.foreco.2013.03.031.

Eyre AW, Wang M, Oh Y, Dean RA. 2019. Identification and characterization of the core rice seed microbiome. Phytobiomes J. 3:148–157.

Fukami T. 2015. Historical contingency in community assembly: Integrating niches, species pools, and priority effects. Annu Rev Ecol Evol Syst. 46:1–23, doi:10.1146/annurev-ecolsys-110411-160340.

Galanti L, Shasha D, Gunsalus K. 2017. Pheniqs: Fast and flexible quality-aware sequence demultiplexing. BioRxiv.

Gardes M, Bruns TD. 1993. ITS primers with enhanced specificity for basidiomycetes - application to the identification of mycorrhizae and rusts. Mol Ecol. 2:113–118.

Gernandt DS, Liston A. 1999. Internal transcribed spacer region evolution in Larix and Pseudotsuga (Pinaceae). Am J Bot. 86:711–723, doi:10.2307/2656581.

Gómez-Gallego M, Leboldus J, Bader MK-F, Hansen E, Donaldson L, Williams NM. 2019. Contrasting the pathogen loads in co-existing populations of Phytophthora pluvialis and Nothophaeocryptopus gaeumannii in Douglas-fir plantations in New Zealand and the US Pacific Northwest. Phytopathology. doi:10.1094/PHYTO-12-18-0479-R.

Grady KL, Sorensen JW, Stopnisek N, Guittar J, Shade A. 2019. Assembly and seasonality of core phyllosphere microbiota on perennial biofuel crops. Nat Commun. 10:4135, doi:10.1038/s41467-019-11974-4.

Hamady M, Walker JJ, Harris JK, Gold NJ, Knight R. 2008. Error-correcting barcoded primers allow hundreds of samples to be pyrosequenced in multiplex. Nat Methods. 5:235–237, doi:10.1038/nmeth.1184.

Hoefnagels MH, Linderman RG. 1999. Biological suppression of seedborne Fusarium spp. during cold stratification of Douglas fir seeds. Plant Dis. 83:845–852.

Hoff JA, Klopfenstein NB, McDonald GI, Tonn JR, Kim M-S, Zambino PJ, Hessburg PF, Rogers JD, Peever TL, Carris LM. 2004. Fungal endophytes in woody roots of Douglas-fir (Pseudotsuga menziesii) and ponderosa pine (Pinus ponderosa). For Pathol. 34:255–271.

Huse SM, Ye Y, Zhou Y, Fodor AA. 2012. A Core Human Microbiome as Viewed through 16S rRNA Sequence Clusters. PLoS ONE. 7:e34242, doi:10.1371/journal.pone.0034242.

James RL, Dumroese RK, Gilligan CJ, Wenny DL. 1989. Pathogenicity of Fusarium isolates from Douglas-fir seed and container-grown seedlings. Pathog Fusarium Isol Douglas-Fir Seed Contain-Grown Seedl.

Klaedtke S, Jacques M-A, Raggi L, Préveaux A, Bonneau S, Negri V, Chable V, Barret M. 2016. Terroir is a key driver of seed-associated microbial assemblages. Environ Microbiol. 18:1792–1804, doi:10.1111/1462-2920.12977.

Kõljalg U, Larsson K-H, Abarenkov K, Nilsson RH, Alexander IJ, Eberhardt U, Erland S, Høiland K, Kjøller R, Larsson E, Pennanen T, Sen R, Taylor AFS, Tedersoo L, Vrålstad T. 2005. UNITE: a database providing web-based methods for the molecular identification of ectomycorrhizal fungi. New Phytol. 166:1063–1068, doi:10.1111/j.1469-8137.2005.01376.x.

Leibold MA, Chase JM. 2018. Metacommunity Ecology. Princeton University Press.

Martin M. 2011. Cutadapt removes adapter sequences from high-throughput sequencing reads. EMBnet. 17.

McMurdie PJ, Holmes S. 2013. phyloseq: An R Package for Reproducible Interactive Analysis and Graphics of Microbiome Census Data. PLOS ONE. 8:e61217, doi:10.1371/journal.pone.0061217.

Morgenstern K, Döring M, Krabel D. 2014. Rhabdocline needle cast◻—◻most recent findings of the occurrence of Rhabdocline pseudotsugae in Douglas-fir seeds. Botany. 92:465–469, doi:10.1139/cjb-2013-0238.

Mujic AB. 2015. Symbiosis in the Pacific Ring of Fire◻: evolutionary-biology of Rhizopogon subgenus Villosuli as mutualists of Pseudotsuga [doctoral dissertation].

Nelson EB. 2018. The seed microbiome: Origins, interactions, and impacts. Plant Soil. 422:7–34, doi:10.1007/s11104-017-3289-7.

Newcombe G, Harding A, Ridout M, Busby PE. 2018. A Hypothetical Bottleneck in the Plant Microbiome. Front Microbiol. 9, doi:10.3389/fmicb.2018.01645.

Okansen J, Blanchet FG, Friendly M, Kindt R, Legendre P, McGlinn D, Minchin PR, O’Hara RB, Simpson GL, Solymos P, Stevens MHH, Szoecs E, Wagner H. 2019. Community Ecology Package.

Proseed NZ. 2019. Douglas-fir.

Pruim R, Kaplan DT, Horton NJ. 2019. mosaic: Project MOSAIC Statistics and Mathematics Teaching Utilities.

Raghavendra AKH, Newcombe G, Shipunov A, Baynes M, Tank D. 2013. Exclusionary interactions among diverse fungi infecting developing seeds of Centaurea stoebe. FEMS Microbiol Ecol. 84:143–153, doi:10.1111/1574-6941.12045.

Rezki S, Campion C, Simoneau P, Jacques M-A, Shade A, Barret M. 2018. Assembly of seed-associated microbial communities within and across successive plant generations. Plant Soil. 422:67–79, doi:10.1007/s11104-017-3451-2.

Rodriguez RJ, White Jr JF, Arnold AE, Redman RS. 2009. Fungal endophytes: diversity and functional roles. New Phytol. 182:314–330, doi:10.1111/j.1469-8137.2009.02773.x.

Rognes T, Flouri T, Nichols B, Quince C, Mahé F. 2016. VSEARCH: a versatile open source tool for metagenomics. PeerJ. 4:e2584, doi:10.7717/peerj.2584.

Salisbury PJ. 1955. Molds of stored Douglas-fir seed in British Columbia. Victoria, British Columbia: Forest Biology Laboratory.

Schoch CL, Seifert KA, Huhndorf S, Robert V, Spouge JL, Levesque CA, Chen W, Fungal Barcoding Consortium, Fungal Barcoding Consortium Author List, Bolchacova E, Voigt K, Crous PW, Miller AN, Wingfield MJ, Aime MC, An K-D, Bai F-Y, Barreto RW, Begerow D, Bergeron M-J, Blackwell M, Boekhout T, Bogale M, Boonyuen N, Burgaz AR, Buyck B, Cai L, Cai Q, Cardinali G, et al. 2012. Nuclear ribosomal internal transcribed spacer (ITS) region as a universal DNA barcode marker for Fungi. Proc Natl Acad Sci. 109:6241–6246, doi:10.1073/pnas.1117018109.

Shade A, Handelsman J. 2012. Beyond the Venn diagram: the hunt for a core microbiome. Environ Microbiol. 14:4–12, doi:10.1111/j.1462-2920.2011.02585.x.

Shade A, Stopnisek N. 2019. Abundance-occupancy distributions to prioritize plant core microbiome membership. Curr Opin Microbiol. 49:50–58, doi:10.1016/j.mib.2019.09.008.

Sherwood-Pike M, Stone JK, Carroll GC. 1986. Rhabdocline parkeri, a ubiquitous foliar endophyte of Douglas-fir. Can J Bot. 64:1849–1855, doi:10.1139/b86-245.

Stamp N. 2003. Out Of The Quagmire Of Plant Defense Hypotheses. Q Rev Biol. 78:23–55, doi:10.1086/367580.

Stopnisek N, Shade A. 2019. Discovery of a spatially and temporally persistent core microbiome of the common bean rhizosphere. Microbiology.

Stucky BJ. 2012. SeqTrace: A Graphical Tool for Rapidly Processing DNA Sequencing Chromatograms. J Biomol Tech JBT. 23:90–93, doi:10.7171/jbt.12-2303-004.

Sturm M, Schroeder C, Bauer P. 2016. SeqPurge: highly-sensitive adapter trimming for paired-end NGS data. BMC Bioinformatics. 17:208, doi:10.1186/s12859-016-1069-7.

Timm CM, Carter KR, Carrell AA, Jun S-R, Jawdy SS, Vélez JM, Gunter LE, Yang Z, Nookaew I, Engle NL, Lu T-YS, Schadt CW, Tschaplinski TJ, Doktycz MJ, Tuskan GA, Pelletier DA, Weston DJ. 2018. Abiotic Stresses Shift Belowground Populus-Associated Bacteria Toward a Core Stress Microbiome. MSystems. 3, doi:10.1128/mSystems.00070-17.

Toju H, Tanabe AS, Yamamoto S, Sato H. 2012. High-Coverage ITS Primers for the DNA-Based Identification of Ascomycetes and Basidiomycetes in Environmental Samples. PLOS ONE. 7:e40863, doi:10.1371/journal.pone.0040863.

Turnbaugh PJ, Hamady M, Yatsunenko T, Cantarel BL, Duncan A, Ley RE, Sogin ML, Jones WJ, Roe BA, Affourtit JP, Egholm M, Henrissat B, Heath AC, Knight R, Gordon JI. 2009. A core gut microbiome in obese and lean twins. Nature. 457:480–484, doi:10.1038/nature07540.

Uchytil RJ. 1991. Pseudotsuga menziesii var. menziesii.

Vandenkoornhuyse P, Quaiser A, Duhamel M, Van AL, Dufresne A. 2015. The importance of the microbiome of the plant holobiont. New Phytol. 206:1196–1206, doi:10.1111/nph.13312.

Vilgalys R. 2018. Conserved primer sequences for PCR amplification and sequencing from nuclear ribosomal RNA.

Watts A, Bansal S, Harrington C, St. Clair B. 2015. Predicting Douglas-Fir’s Response to a Warming Climate. Sci Find. 1–6.

White TJ, Bruns T, Lee S, Taylor JL. 1990. Amplification and direct sequencing of fungal ribosomal RNA genes for phylogenetics. PCR Protoc Guide Methods Appl. 18:315–322.

Wickham H. 2016. ggplot2: Elegant Graphics for Data Analysis. Springer. 266 p.

Wright E S. 2016. Using DECIPHER v2.0 to Analyze Big Biological Sequence Data in R. R J. 8:352, doi:10.32614/RJ-2016-025.

